# The causal effect of educational attainment on Alzheimer’s disease: A two-sample Mendelian randomization study

**DOI:** 10.1101/127993

**Authors:** Emma L Anderson, Kaitlin H Wade, Gibran Hemani, Jack Bowden, Roxanna Korologou-Linden, George Davey Smith, Yoav Ben-Shlomo, Laura D Howe, Evie Stergiakouli

## Abstract

**Background:** Observational evidence suggests that higher educational attainment is protective for Alzheimer’s disease (AD). It is unclear whether this association is causal or confounded by demographic and socioeconomic characteristics. We examined the causal effect of educational attainment on AD in a two-sample MR framework.

**Methods:** We extracted all available effect estimates of the 74 single nucleotide polymorphisms (SNPs) associated with years of schooling from the largest genome-wide association study (GWAS) of educational attainment (N=293,723) and the GWAS of AD conducted by the International Genomics of Alzheimer’s Project (n=17,008 AD cases and 37,154 controls). SNP-exposure and SNP-outcome coefficients were combined using an inverse variance weighted approach, providing an estimate of the causal effect of each SD increase in years of schooling on AD. We also performed appropriate sensitivity analyses examining the robustness of causal effect estimates to the various assumptions and conducted simulation analyses to examine potential survival bias of MR analyses.

**Findings:** With each SD increase in years of schooling (3.51 years), the odds of AD were, on average, reduced by approximately one third (odds ratio= 0.63, 95% confidence interval [CI]: 0.48 to 0.83, p<0.001). Causal effect estimates were consistent when using causal methods with varying MR assumptions or different sets of SNPs for educational attainment, lending confidence to the magnitude and direction of effect in our main findings. There was also no evidence of survival bias in our study.

**Interpretation:** Our findings support a causal role of educational attainment on AD, whereby an additional ∼3.5 years of schooling reduces the odds of AD by approximately one third.

## INTRODUCTION

Observational evidence for modifiable risk factors for Alzheimer’s disease (AD) is inconsistent, with studies reporting positive, negative and null findings for the same risk factor.(1) Education, however, is an exception with the majority of studies reporting higher levels of education to be associated with a reduction in both incidence and prevalence of dementia, in various age, social and ethnic groups.(2, 3) There has been a continued debate as to whether the association between education and AD is likely to be causal, or confounded by other demographic and socioeconomic characteristics that are associated with both educational attainment and risk of AD.(4-6) Establishing whether or not a potential risk factor is causally related to a disease ensures that prevention efforts are appropriately targeted.

Observational epidemiological studies are prone to biases such as confounding and reverse causation.(7) Mendelian randomization (MR) is a method that uses genetic variants to proxy for environmental exposures, to avoid such bias and to generate more reliable causal estimates.(8) Two existing studies (9, 10) have investigated whether educational attainment has a causal effect on AD using MR.(11) Both studies used one single nucleotide polymorphisms (SNPs) that was previously identified as being associated with the number of years of education, and two SNPs identified as being associated with the probability of completing university, in a large genome-wide association study (GWAS, n=101,069).(12) However, the findings are conflicting: the first found little evidence of an association between university completion and AD (OR: 0.95, 95% CI: 0.67–1.34), and weak evidence of an association between years of schooling (OR: 0.71, 95% CI: 0.48– 1.06) and AD.(10) The second reported an average 1.1% lower dementia risk per year of schooling (95% CI: −2.4 to 0.02%).

A larger (n=293,723) genome-wide association study was published in 2016, which identified 74 loci as being independently associated with years of schooling,(13) and explaining a larger proportion of the variance in years of schooling than the previous GWAS (0.06% compared with 0.02%). The results of this study have not yet been utilized to assess the causal effect of educational attainment on AD. Using more SNPs that are robustly associated with years of schooling in MR analyses to assess whether or not educational attainment has a causal effect on AD, will provide greater statistical power and therefore will improve the evidence base for important question. In this study, we aimed to use the larger list of SNPs associated with years of schooling from this updated GWAS in a two-sample MR framework. We also aimed to improve on the previous MR studies of educational attainment and AD by conducting a comprehensive series of sensitivity analyses, to establish whether the observed causal effect is robust to the various assumptions and potential biases of MR analyses.(7)

## ANALYSES

All analyses were conducted in MR-Base, which includes complete summary data from 1094 GWAS on diseases and other complex traits into a centralised database and an analytical platform that uses these data to perform MR tests and sensitivity analyses. (www.mrbase.org).(14) In our study, MR-base was employed to perform a two-sample MR analysis. Two-sample MR provides an estimate for the causal effect of an exposure on an outcome, using summary results from two independent genome-wide association studies.(7)

### Samples

The most recent GWAS of educational attainment (n=293,723)(13) identified 74 approximately independent genome-wide significant (p<5x10^-8^) loci that were associated with years of schooling in the discovery sample. For each locus, the ‘lead SNP’ was defined as the SNP in each independent genomic region with the smallest p-value. These 74 SNPs explain 0.06% of phenotypic variation in years of schooling. We extracted the effect estimates and corresponding standard errors of the 74 SNPs from the educational attainment GWAS (13) and the large-scale GWAS of AD conducted by the International Genomics of Alzheimer’s Project (IGAP) (n=17,008 AD cases and 37,154 controls).(15) Of the 74 SNPs, three were not present in the AD GWAS study (and no SNPs in linkage disequilibrium (LD) were available) and eight palindromic SNPs were excluded at the harmonisation stage (i.e. aligning effect alleles from both exposure and outcome datasets) (16) due to missing effect allele frequencies (Figure 1).(16) Thus, a total of 63 SNPs were used for the main analysis to investigate the association between years of schooling and AD.

**Figure 1:**
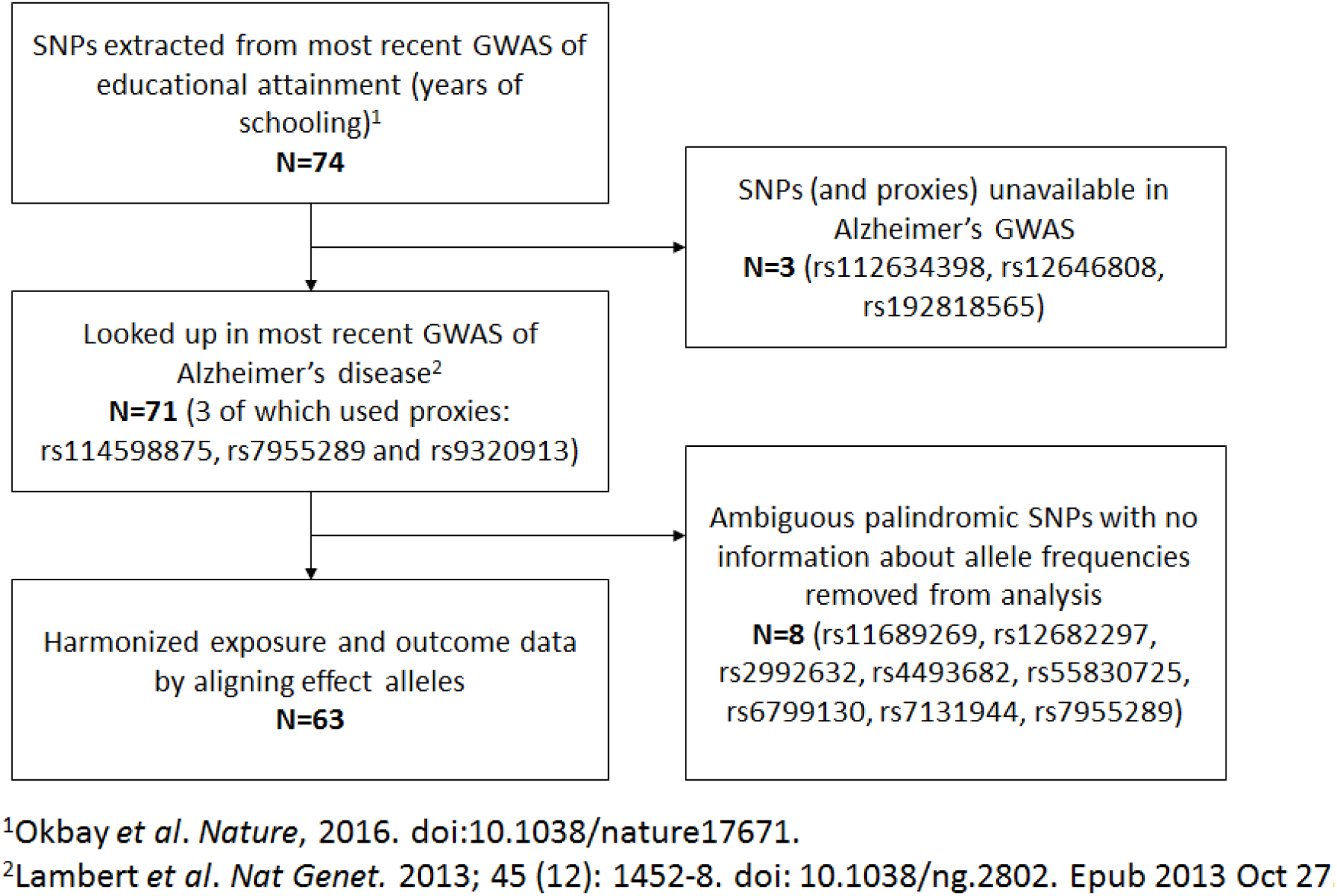
MR Analysis methods

### Estimating the causal effect of years of schooling on Alzheimer’s disease

The SNP-exposure (years of schooling, in standard deviation (SD) units) and SNP-outcome (AD, log odds ratios [ORs]) coefficients were combined using an inverse-variance-weighted (IVW) approach to give an overall estimate of the causal effect across all SNPs. This is equivalent to a weighted regression of the SNP-outcome coefficients on the SNP-exposure coefficients with the intercept constrained to zero (i.e. no pleiotropy). This method assumes the gene-exposure association estimates are measured without error (the no measurement error [NOME] assumption).(17) The results of the analyses were converted to ORs, representing the odds of AD per one SD (one SD=3.51 years) increase in years of schooling.

### Sensitivity analyses

A number of sensitivity analyses were conducted to interrogate the robustness of our findings and validity of genetic instruments used. We compared results from our main analyses to those obtained with MR-Egger regression,(17) as the use of multiple alleles in MR analyses increases the potential for pleiotropic effects due to aggregation of invalid genetic instruments.(16) MR-Egger assumes NOME but relaxes the assumption that the effects of genetic variants on the outcome operate entirely via the exposure, by not constraining the intercept term to zero in the weighted regression described above. In this instance, the intercept parameter indicates the overall pleiotropic effect of the SNPs on the outcome (i.e. a direct effect on the outcome independent of the exposure, which would violate MR assumptions), with a non-zero intercept providing evidence for bias in the causal estimate due to pleiotropy. MR-Egger estimates remain consistent only if the magnitude of the gene exposure associations across all variants are independent of their pleiotropic effects (i.e. the InSIDE assumption).(17) The beta coefficient (or slope) of MR-Egger provides a causal estimate of the exposure on the outcome, accounting for this level of pleiotropy and assuming that the pleiotropic effect of SNPs on the outcome is not correlated with the instrument strength.(17) As recommended by Bowden et al. (9), the extent to which pleiotropy was balanced across the set of instruments was visually assessed by plotting the causal effect estimates against their precision, using a funnel plot and checking for asymmetry. We assessed the NOME assumption using an adaptation of the I^2^ statistic (18) to the two-sample summary data MR context, which is referred to as I^2^_GX_. This sensitivity analysis accounts for uncertainty in the SNP-exposure estimates. I^2^_GX_ provides an estimate of the degree of regression dilution in the MR-Egger causal estimate. We then used simulation extrapolation (SIMEX) to adjust the MR-Egger estimate for dilution, as described in (19).

We compared results from our main analyses to those obtained with the weighted median method.(20) The weighted median method provides a consistent estimate of causal effect if at least 50% of the weight in the analysis stems from genetic variants are valid instrumental variables for the exposure (i.e. robustly associated with the exposure, not associated with confounding factors and only associated with the outcome via the exposure of interest).

We conducted a leave-one-out permutation in which each SNP was systematically removed from the analysis, to assess the influence of potentially pleiotropic SNPs on the causal estimates.(21)

Given that we present our main IVW, MR-Egger and weighted median results using the 63 SNPs published on the discovery sample of the educational attainment GWAS, we conducted a series of sensitivity analysis to check whether our results were sensitive to ‘winner’s curse’ (i.e. where the effect sizes of variants identified within a single sample are, likely to be larger than in the overall population, even if they are truly associated with the exposure). We did this by investigating how sensitive our causal estimates are to using (i) different sets of SNPs as instrumental variables obtained from replication samples in UK Biobank and (ii) different SNP effect size estimates for years of schooling (Supplemental Table A).

It is known that the estimated effect of an exposure on an outcome, that are both associated with mortality (and thereby, study participation), may be susceptible to survival bias.(22) This may also potentially be a problem for two-sample MR. For example, if individuals with lower educational attainment are more likely to die before the age of onset of AD, bias may occur because those individuals with a genetic predisposition for higher educational attainment are likely to live longer, thus having greater risk of being diagnosed with AD. This phenomenon, commonly referred to as collider bias in epidemiology,(23) could induce a nonzero causal effect estimate, whereby higher educational attainment appears to be associated with greater risk of AD, even if no true biological association exists. To investigate whether our causal estimates may be biased by survival, we used methods that have previously been applied to assess the impact of survival bias in an MR study of body mass index and Parkinson’s disease.( Noyce et al, 2017; in press) Simulations were performed, where a large sample (n=500,000) was generated with data on: (i) educational attainment, (ii) SNPs associated with educational attainment, (iii) age, (iv) mortality status, and (vi) AD status. Observational and MR analyses were performed on the simulated data to gauge the extent to which an association between educational attainment and AD was induced artificially by survival bias. The entire process was repeated 1000 times to obtain a distribution of effect sizes due to frailty effects only. Full details of method are outlined in Noyce et al (Noyce et al, 2017; in press) and details of our own analyses are provided in the online supplement. All analyses were undertaken in R (version 3·2·3).

## RESULTS

### Estimating the causal effect of years of schooling on Alzheimer’s disease

There was strong evidence to suggest that years of schooling have a causal effect on AD. With each SD increase in years of schooling (3.51 years), the odds of AD were, on average, reduced by approximately one third (OR= 0.63, 95% confidence interval [CI]: 0.48 to 0.83, p<0.001, Figure 2, Table 1 and Supplemental Figure A).

**Figure 2:**
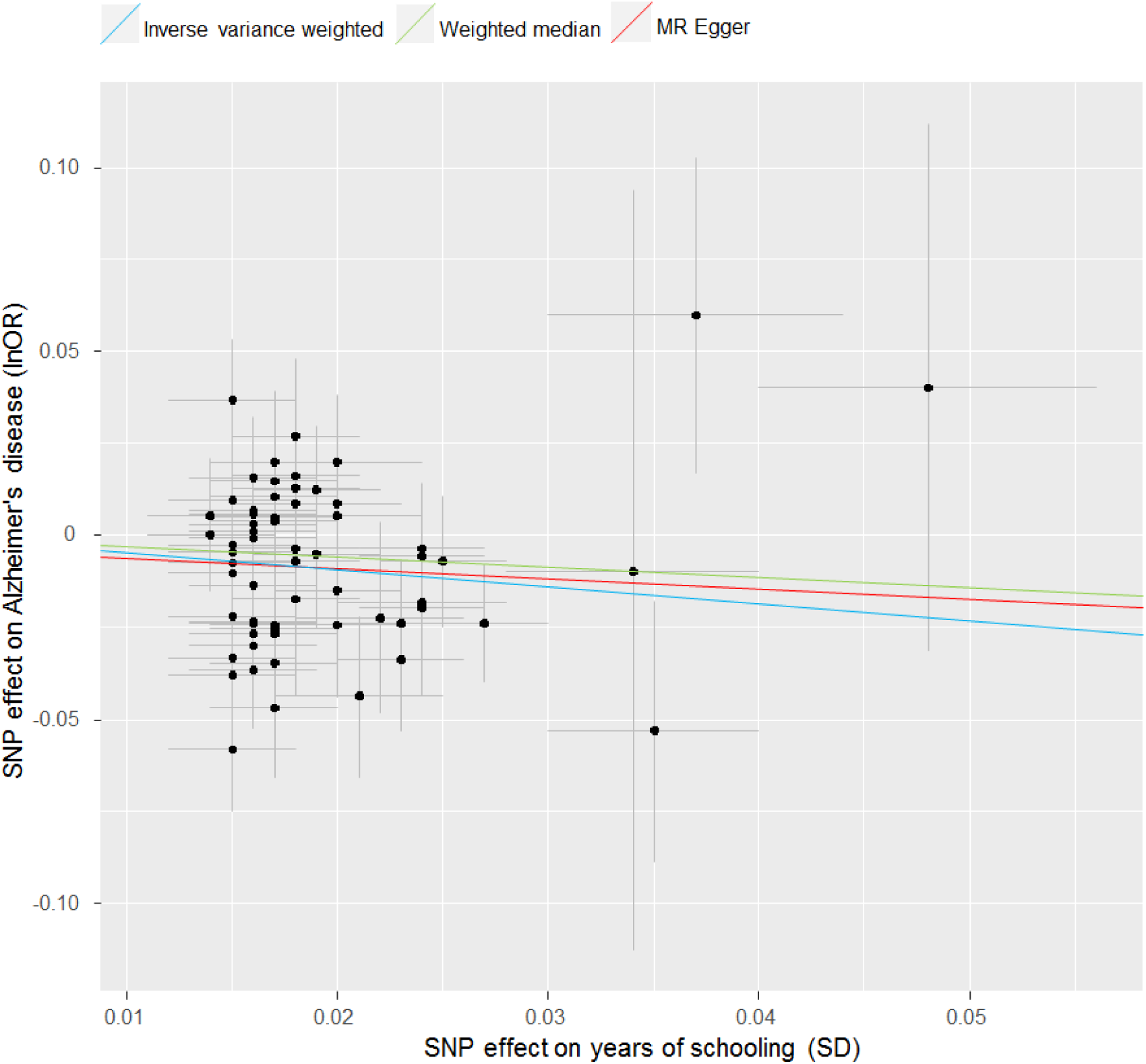
Comparison of results for each different MR method

**Table 1:** Causal effect of years of schooling on AD in each MR method, using 63 SNPs from the discover GWAS of educational attainment

| **MR analysis** | **OR** | **95% CI** | **P** |
| --- | --- | --- | --- |
| **IVW** | 0.63 | 0.48 to 0.83 | <0.001 |
| **Weighted Median** | 0.75 | 0.53 to 1.08 | 0.12 |
| **MR-Egger** | 0.72 | 0.18 to 2.92 | 0.64 |

### Sensitivity analyses

MR-Egger regression indicated little evidence of pleiotropy (intercept: -0.003, 95% CI: -0.03 to 0.02, p=0.79) and causal effect estimates were very similar to our main analyses (odds ratio for AD per 1 SD higher years of education = 0.72 (95% CI: 0.18 to 2.92), Figure 2, Table 1); however, the standard error was expectedly much larger, because MR-Egger estimates two parameters (i.e. both an intercept and a slope) compared to the single parameter that both IVW and weighted median methods estimate (i.e. only the slope). Assessment of the NOME assumption with respect to the MR-Egger estimate gave I^2^_GX_ = 0.67, suggesting an expected 33% attenuation of the causal estimate toward zero, due uncertainty in the SNP-exposure exposure estimates. Bias adjustment via SIMEX gave a corrected MR-Egger causal estimate of 0.63 (95% CI: 0.08, 5.13) for AD per SD (3.5 years) increase in years of schooling, which is consistent with the IWV effect estimate.

The weighted median estimate was slightly attenuated from the IVW estimate (odds of AD were, on average, 0.75 (95% CI: 0.53 to 1.08, p=0.13) per SD increase in years of schooling, Figure 2, Table 1).

Compared to the main IVW analysis, the causal effect estimates were very similar after removing one SNP over any other (i.e. the leave one out permutations, Supplemental Figure B). Had any one of the SNPs been invalid, it is likely we would have observed some distortion in the distribution of results in these analyses.

Causal effect estimates from the IVW, MR-Egger and weighted median regression analyses were slightly larger when using the SNP effect sizes estimated in the UK Biobank replication sample (n=111,349, OR=0.54, 95% CI: 0.4-0.72, P=<0.01). This is consistent with winner’s curse diluting the causal effect estimate when using SNP effect sizes from the discovery sample (as in our main analysis). This new causal effect estimate was not sensitive to using different replication thresholds (Supplemental Table A). We also performed the analysis again using SNPs that were only discovered in UK Biobank (not in the discovery sample) and had not previously been used as instruments, and the effect was in the same direction as the main analysis.

Results from our null model simulations suggested that survival bias could yield biased causal estimates in observational studies (p < 0.01, Table A of the online supplement), but induced no substantive effect in this particular MR study (p = 0.91, Table 1 of the online supplement), meaning that survival bias is not likely to be influencing the estimate of the causal effect.

## DISCUSSION

Our study assessed whether educational attainment has a causal effect on risk of AD, using 63 genetic variants identified in the most recent educational attainment GWAS in a two-sample MR framework. Our findings suggest that, per additional 3.5 years of schooling, odds of AD reduce by 25-37%. Estimates were relatively consistent across each of the three different MR methods (IVW, weighted median and MR-Egger), which all have varying assumptions. The IVW makes the strong assumption that all genetic variants used are valid instrumental variables, whereas the weighted median method provides a consistent estimate of causal effect if at least 50% of the information in the analysis comes from variants that are valid instruments. The causal estimate from the weighted median regression was slightly attenuated from the IVW, which may suggest some of the genetic variants are not valid instruments. Nevertheless, the effect estimate was consistent with the IVW and results were robust to the removal of each SNP in turn in the leave-one-out permutations. Furthermore, no pleiotropy was detected within the set of 63 genetic variants used, as reflected by the estimate of the MR-Egger intercept, and the causal estimate from MR-Egger was similar to that from the IVW regression and weighted median method. MR-Egger confidence intervals were much wider, but that is to be expected: It fits a linear regression model using the SNP-exposure association estimates as the explanatory variable, in order to check for agreement in causal estimates across genetic instruments. As with any regression model that estimates an intercept and a slope, a loss in precision will always occur relative to the same regression model but with no intercept (e.g. IVW), especially when there is little variation in the explanatory variable (e.g. the SNP-exposure associations). Assessment of the NOME assumption suggests an attenuation of the MR-Egger estimate by 33%, due to uncertainty in the SNP-exposure estimates. When the MR-egger causal effect estimate is adjusted for this regression dilution bias, the effect estimate is consistent with the IVW effect estimate.

Our findings are consistent with the ‘brain reserve’ and the ‘cognitive reserve’ hypotheses. Brain refers to structural differences in the brain itself that may increase tolerance of pathology, whereas cognitive reserve refers to differences in the ability to tolerate and compensate for the effects of brain atrophy, using pre-existing cognitive-processing approaches or compensatory mechanisms.(24) In support of this, higher levels of education have been shown to increase whole brain and ventricular volume as well as cortical thickness.(25-27) A post-mortem study of 130 elderly patients who had undergone cognitive assessment approximately 8 months before death also showed that, at any given level of brain pathology, higher education was associated with better cognitive function.(28) In addition, higher educational attainment may lead to extrinsic compensation through adaptations.

Hence, more educated people will usually have occupations that are more intellectually demanding or have greater resources to partake in intellectual activities, resulting in greater cognitive stimulation and consistent with the “use it or lose it” hypothesis.(6) Together, these studies and ours suggest that the mechanism through which higher educational attainment confers protection against advancing AD pathology may, at least in part, be driven by the increased time it takes for an individual to reach the threshold of cognitive decline, whereby functional impairment starts to adversely impact on daily living and a clinical AD diagnosis is made.

Our findings suggest that the relatively recent change in school policy in the United Kingdom (in 2013), which now requires young people to remain in at least part-time education until age 18 years (as opposed to 16 years), may have beneficial consequences in for AD risk in the future. It is worth noting that the educational attainment GWAS only assessed years of full-time academic training from primary education through to advanced research qualifications (e.g. PhD). Therefore, it remains unclear whether the same genetic variants would be associated with other aspects of education, such as completing vocational courses or completing part-time as opposed to full-time courses. It’s not clear whether education needs to be completed in a formal setting (such as school or college), or whether any form of learning (e.g. learning new skills ‘on the job’ such as in an apprenticeship during adolescence, or through career development and training courses as an adult in existing fulltime employment) would confer the same degree of cognitive protection. This likely depends on the mechanism driving the association between education and AD, thus further studies to unpick the mechanisms may help to shed light on which forms of learning may confer cognitive benefits later in life. Finally, it is not clear whether the cognitive benefit would still be apparent in those people who left school at 16 to work full time, but returned to education (either full-time or part-time) later in life (e.g. in mid-life).

### Comparison to other studies

Our findings extend upon an extremely large and diverse (i.e. in different age, ethnic and socioeconomic groups) body of observational evidence suggesting higher educational attainment is associated with lower risk of AD.(2, 3) A meta-analysis of 69 different studies reported a pooled OR for any dementia for individuals with low education level compared to with high education of 2.61 (95% CI 2.21–3.07, p<0.001). Our findings also support the two previous MR studies that have been conducted to assess a possible causal role of educational attainment on risk of AD.(9, 10) The first study used three SNPs identified in the previous GWAS of educational attainment in a two-sample MR framework; however, two of these SNPs were identified within the context of completion of university (a binary variable) and one SNP was associated with number of years of schooling. Combining these three SNPs into a single genetic score and using this as an instrumental variable for years of schooling, the authors reported weak evidence of an inverse association between years of schooling and AD risk (OR: 0.71, 95% CI: 0.48– 1.06, p=0.097). Given that education is a complex and heterogeneous phenotype, it may not be robustly explained by just three SNPs; those 3 SNPs explain approximately 0.02% of the variance in educational attainment (as opposed to 0.06% explained by the 74 SNPs in the most recent GWAS), which may also explain why the strength of evidence was weaker in that study than in ours. The second study was a one-sample MR analysis conducted in the Health and Retirement Study cohort. The authors used two separate instruments: (1) a genetic risk score for educational attainment based on three SNPs and (2) compulsory schooling laws and state school characteristics (term length, student teacher ratios, and expenditures). Their results showed a 1.1% reduction in dementia risk per year of additional schooling and an even larger protective effect when using compulsory schooling laws and state school characteristics as instrumental variables. Thus, each of the MR studies to date support educational attainment having a causal effect on AD risk. Finally, using the same GWAS of education, an inverse genetic correlation was reported between educational attainment and AD (R: -0.29, standard error: 0.1, p=5×10^−3^), which is consistent with our findings.(29)

### Strengths and limitations

Our study used the most comprehensive set of SNPs identified to date from the most recent GWAS of educational attainment to assess whether educational attainment has a causal effect on AD. MR studies are comparable to randomized controlled trials, provided all assumptions are valid. Thus, bias by confounding (for example by lifestyle and environmental factors) and reverse causation are unlikely. The first assumption of MR is that the genetic instruments are associated with the exposure, and this is likely valid given the SNPs used in our analyses were identified in the largest educational attainment GWAS to date. The second assumption of MR is that the possible confounders of the association between educational attainment and AD are not associated with the genetic instruments. We are unable to formally test this using summary data in the two-sample MR framework, however, there is empirical evidence that this is unlikely to be the case.(30) The third assumption is that the genetic instruments are not associated with the outcome other than through the exposure variable, and this is likely to be valid given our MR-Egger intercept estimates showed no evidence of pleiotropy.

In two-sample MR, “winner’s curse” (the notion that effect sizes of variants identified within a single sample are likely to be larger than in the overall population, even if they are truly associated with the exposure) can bias causal estimates if two overlapping samples are used.(31) This is not the case in the current investigation, as there is no overlap between cohorts used in the educational attainment GWAS and the AD GWAS. Winner’s curse can also bias causal estimates if genetic variants from the discovery phase of the exposure GWAS are used. In our study, causal estimates were consistent even when using only the seven SNPs that replicated at genome-wide significance in the educational attainment GWAS. Furthermore, winner’s curse is likely to bias causal effect estimates towards the null and not to the observational estimate.

## Conclusions

Findings from our MR study support a causal role of educational attainment on AD. A range of sensitivity analyses produced consistent results, lending more confidence to the causal estimates and direction of effect (i.e. higher educational attainment reducing odds of AD), and suggesting that our estimates are not largely driven by horizontal pleiotropy. Further studies should investigate whether the magnitude of effect of educational attainment on AD is similar in people who complete academic vs vocational training, part-time vs full-time, and early life vs later-life education.

